# Morphology-robust quantification of subcellular organization in complex cells

**DOI:** 10.64898/2026.05.28.728543

**Authors:** Robert Hu, Nima N. Naseri, Ophir Shalem, Pablo G. Camara

## Abstract

Quantitative analysis of subcellular protein organization is often confounded by variation in cell morphology, limiting the identification and interpretation of localization patterns in fluorescence microscopy data from morphologically complex cells, such as neurons and glia. We introduce CellAligner, an unsupervised framework that uses fused unbalanced Gromov-Wasserstein couplings to map protein distributions from morphologically distinct cells into shared anchor-cell geometries, enabling morphology-robust comparison of subcellular localization. In neuronal imaging benchmarks, applying current image-analysis methods (CellProfiler, Cytoself, Paired Cell Inpainting) to CellAligner’s anchor-cell representations substantially reduced morphology-associated confounding while approximately doubling their multiclass MCC for localization classification. We demonstrate its biological utility by identifying U18666A-induced lysosomal trafficking defects in human iPSC-derived neurons. To scale the approach, we developed dCellAligner-OT, a fast deep metric learning model that approximates CellAligner’s optimal transport distances and anchor-cell representations, enabling atlas-scale analyses. CellAligner provides a general framework for morphology-robust analysis of subcellular organization in complex cellular systems.

## Introduction

The precise spatial localization of proteins within cells is fundamental to cellular function and human health, as it controls protein interactions, post-translational modifications, and signaling cascades^1^. Protein mislocalization is mechanistically implicated in a wide range of human diseases, with mutations in over 300 intracellular trafficking genes linked to more than 200 disorders^2^, including cancer, neurodegenerative, immune, and cardiovascular diseases. Critically, mislocalization often cannot be inferred from sequence or expression data alone, requiring direct, quantitative measurement of protein distributions within intact cells.

Recent advances in image-based protein profiling and high-resolution fluorescence microscopy have greatly expanded the ability to profile subcellular organization with high multiplexity and submicron resolution^3,4^, yielding increasingly comprehensive maps of subcellular proteome organization^5–7^. In parallel, CRISPR-engineered iPSC reporter collections have enabled systematic visualization of subcellular compartments in iPSC-derived cells and organoids^8,9^. Collectively, these innovations are fueling a new era of image-based cell phenotyping. Yet quantitative characterization of localization from these data remains a major challenge, particularly in cells with complex and heterogeneous morphologies, such as neurons, glia, and primary cells.

Traditional feature-based methods for cell image analysis, such as CellProfiler^10^, rely on engineered descriptors such as intensity statistics and Zernike moments^11^, and are highly effective at capturing morphological and textural properties of cells. These methods have proven very useful in high-content imaging screens^12^. However, they have limited capacity to disentangle subcellular localization from morphological variation, introducing morphology-dependent biases that can compromise the accuracy and interpretability of subcellular localization measurements, particularly in heterogeneous or highly structured cell types.

More recently, self-supervised deep learning methods, such as Paired Cell Inpainting^13^ and Cytoself^14^, have emerged as a powerful paradigm for the analysis of subcellular organization. These methods aim to learn representations that decouple morphological variation from subcellular localization differences. However, achieving robust morphology-invariant representations typically requires large training datasets spanning many proteins and tens of thousands of images, such as the Human Protein Atlas^5^ or OpenCell^6^. As a result, these methods may not reliably generalize beyond the cell types, morphologies, and imaging conditions on which they were trained, facing practical limitations when large-scale training data are unavailable.

Consequently, the field still lacks a general quantitative framework for comparing subcellular protein organization from imaging data in a manner that is robust to morphological differences and does not require large training datasets. While recent work has begun to address this challenge by introducing intracellular coordinate systems based on spherical harmonics^15,16^, these approaches are best suited to approximately spherical or smoothly parameterizable cell geometries and do not provide a general framework for morphology-robust analyses of complex differentiated cell types.

To address this gap, here we develop CellAligner, a general framework for quantifying and comparing subcellular organization in 2D imaging data while explicitly accounting for differences in cell morphology. Gromov-Wasserstein (GW) couplings^17–20^ provide a principled method for building correspondences between cells with different geometries based on internal geometric relationships rather than a shared coordinate system. CellAligner uses these couplings to map protein distributions from morphologically distinct cells into a common cellular geometry, enabling direct comparison of localization patterns using existing quantitative colocalization metrics, such as optimal transport colocalization. Because the correspondences are guided by reference channels, such as nuclear, membrane, and organelle stains, the resulting maps respect compartmental landmarks. The mapped representations can also be analyzed using existing cell image analysis methods, such as CellProfiler, Cytoself, and Paired Cell Inpainting, reducing morphology-dependent biases and improving their ability to distinguish subcellular localization patterns. In neuronal data, this strategy substantially improved the performance of current methods in downstream analyses of subcellular organization. Because CellAligner is unsupervised, it does not require training data and can be applied to small datasets.

To scale this approach to datasets containing millions of cells, we distill CellAligner into a deep metric learning network. The key innovation relative to existing self-supervised approaches is that separation between morphology and subcellular localization is not a hoped-for emergent property of representation learning but a direct consequence of CellAligner supervision. Because the learning signal is derived from CellAligner itself, the resulting model retains the unsupervised, cell-type-agnostic nature of the underlying framework while enabling the speed and scalability required for application to large-scale datasets.

We provide CellAligner and its deep neural network extension in CAJAL^21^, our open-source Python library for metric-geometry-based analysis of cellular morphology, extending its capabilities from morphology to subcellular organization. We expect the morphology-robust analyses enabled by CellAligner will have direct applications to disease mechanism studies, drug screening, and biomarker discovery.

## Results

### Metric geometry enables quantifying subcellular localization and cell morphology independently

The GW coupling between two shapes or *metric spaces* provides a probabilistic mapping that preserves internal geometric relationships^17^. The minimum value of the GW objective defines a morphological distance that satisfies all mathematical properties of a distance metric and is invariant to rotations and translations of the individual shapes. This distance has proven highly useful for quantifying cell morphological variation and constructing summary spaces in which each point represents a cell and pairwise distances represent morphological dissimilarity^21^. In addition, unbalanced and fused extensions of GW^18–20^ enable partial mappings and incorporate additional shape features, such as local fluorescence intensities.

Building upon these concepts, we devised a general, unsupervised method to separate changes in protein subcellular localization from differences in cell morphology (**Fig. 1a**) (see Methods). CellAligner takes as input multi-channel images, including fluorescently labeled proteins and morphological stains (e.g., nuclear or membrane stains), along with cell segmentation masks. To quantify differences in subcellular localization independently of morphology, CellAligner first maps all cells to a shared anchor cell by computing a pixel-wise fused unbalanced GW mapping. This mapping jointly minimizes discrepancies in morphological staining intensities and intracellular distances between each cell and the anchor, and it allows partial matching through unbalanced regularization so that cells with substantially different geometries can be partially mapped. The resulting mapping is used to transfer each cell’s protein distribution onto the morphology of the anchor cell. Differences in subcellular protein localization are then quantified by computing standard colocalization metrics^22–27^ between the mapped protein distributions within the anchor geometry. Here, we use optimal transport (OT) distances, following optimal transport colocalization^27^, due to their natural interpretation as the average intracellular distance over which mass from one protein distribution must be transported to match another. Alternatively, existing cell image analysis methods^10,13,14^ can be applied to the corresponding anchor-cell representations, thereby reducing morphological-associated confounding from downstream analyses.

**Figure 1.**
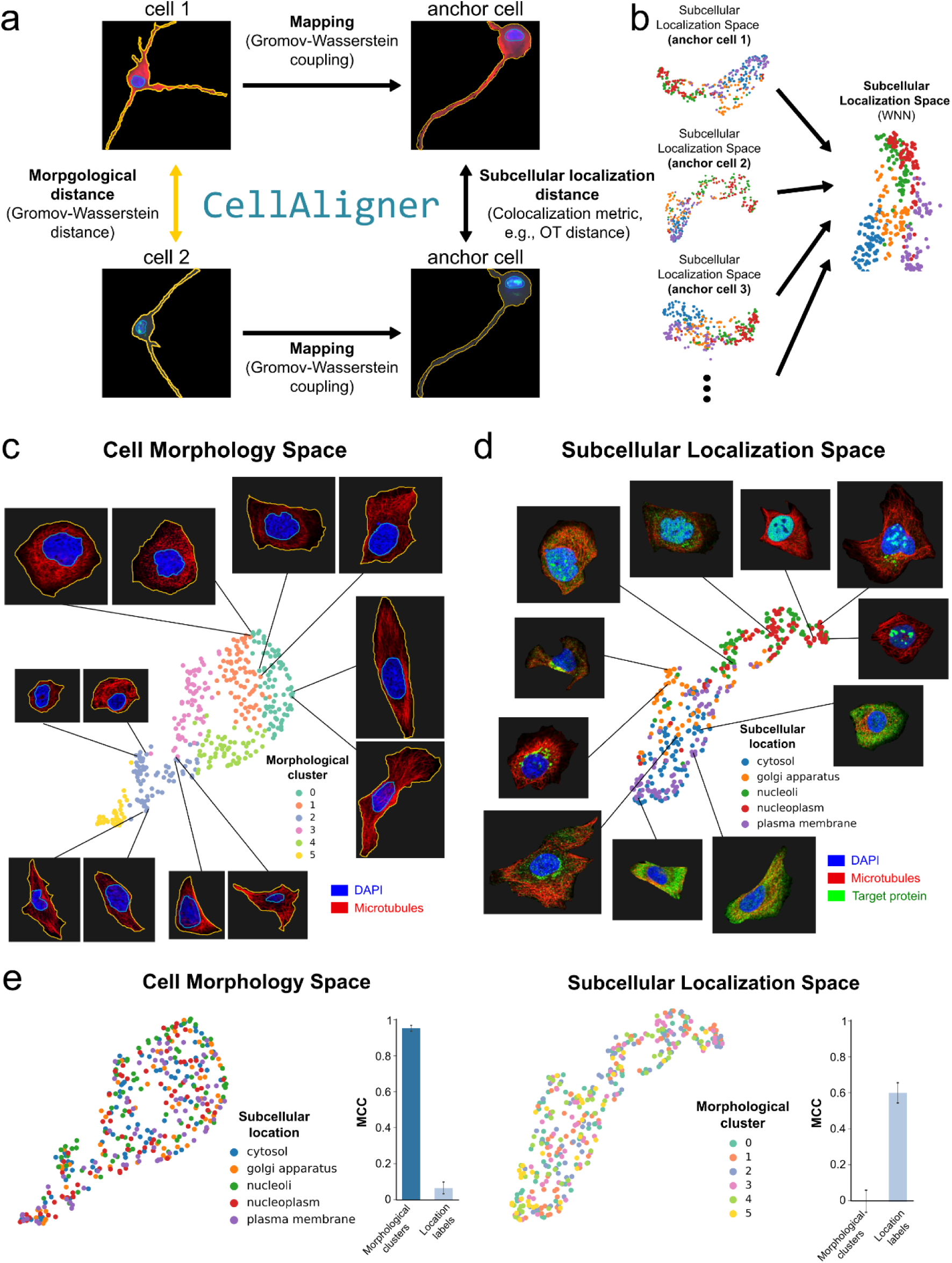
Independent quantification of subcellular localization and cell morphology using metric geometry. **(a, b)** Overview of the CellAligner algorithm. (**a**) Given two query cells and an anchor cell, CellAligner first computes GW couplings that map each query cell onto the anchor cell. It then computes the discrepancy between the mapped protein distributions within the anchor cell using a colocalization metric, such as their optimal transport (OT) distance, yielding a subcellular localization distance. Additionally, CellAligner computes the GW distance between the segmentation masks of the two cells, providing a morphological distance. (**b**) Subcellular localization pairwise distance matrices computed using different anchors are embedded into a high-dimensional Euclidean space and integrated into a unified subcellular localization space using weighted nearest neighbors. **(c-e)** Application of CellAligner-OT to 387 Human Protein Atlas cell images representing 48 proteins. (**c**) UMAP representation of the cell morphology space, colored by the morphological clusters. Representative cell images are shown. (**d**) UMAP representation of the subcellular localization space, colored by the manually annotated subcellular compartment of each protein. (**e**) Left: Cell morphology space colored by the subcellular localization labels and MCC score of a k-nearest neighbor classifier trained to predict subcellular localization labels using the cell morphology space or the subcellular localization space. Right: Subcellular localization space colored by the morphological cluster labels and MCC score of a classifier trained to predict the morphological labels using the cell morphology space or the subcellular localization space.

Repeating the mapping procedure across several morphologically distinct anchors and integrating the resulting distances using a non-coordinate extension of the weighted nearest-neighbor algorithm^28^ yields subcellular localization distances that are robust to differences in cell morphology (**Fig. 1b**) (see Methods). Thus, in addition to a cell morphology summary space, CellAligner produces a subcellular localization summary space in which each point represents a cell and pairwise distances reflect differences in protein localization independent of cell morphology. These localization summary spaces can be integrated across proteins to produce higher-level subcellular organization spaces, in which pairwise distances summarize differences in intracellular organization between cells.

We implemented CellAligner in our open-source Python library CAJAL for the morphometric analysis of cells^21^. To qualitatively assess the ability of CellAligner to decouple subcellular protein localization differences from morphological variation, we first applied it to an immunofluorescence imaging dataset from the Human Protein Atlas^5^, comprising 387 cells and 48 proteins localized across five major subcellular compartments. As anticipated, morphological features, such as cell size, eccentricity, and compactness, appeared regularly distributed in the morphology space (**Fig. 1c** and **Supplementary Fig. 1a**). In contrast, the subcellular localization space effectively captured variation in subcellular protein distribution, with annotated compartment-specific patterns clearly separated in this space (**Fig. 1d**, Matthews Correlation Coefficient (MCC) of a 𝑘 = 5 nearest-neighbor classifier = 0.55, *p*-value < 10^-4^). Crucially, the annotated subcellular localization patterns were interspersed when visualized in the morphology space, whereas morphological features did not form structured patterns in the localization space (**Fig. 1e** and **Supplementary Fig. 1b**), demonstrating that the two CellAligner-derived summary spaces successfully capture distinct and biologically meaningful sources of variation in the imaging data.

### Morphological and image-associated variation confound current subcellular localization representations

To assess whether current methods for subcellular localization analysis are sensitive to unrelated morphological and image-associated variation, we considered three state-of-the-art methods that represent distinct paradigms for subcellular localization analysis. CellProfiler^10^ is an unsupervised, feature-based approach that extracts predefined morphological and textural descriptors such as elongation, radially binned intensities, and Zernike moments, and is widely used in high-content imaging pipelines. On the other hand, Cytoself^14^ and Paired Cell Inpainting^13^ are self-supervised deep learning models trained on the OpenCell^6^ (*n* = 1,048,800 cells) and Human Protein Atlas^5^ (*n* = 638,640 cells) datasets, respectively.

We applied these methods to a Human Protein Atlas immunofluorescence dataset comprising 1,021 annotated cells from 138 images across three cell lines (U-251MG, U2OS, and A-431) (**Fig. 2a**). For each method, we constructed a subcellular localization space from the extracted features or embedding and trained *k*-nearest neighbor classifiers to predict localization label, cell-line identity, and image identity for each cell (**Supplementary Fig. 2**) (see Methods). Performance was evaluated using Matthews correlation coefficient (MCC) under a 10-fold cross-validation scheme.

**Figure 2.**
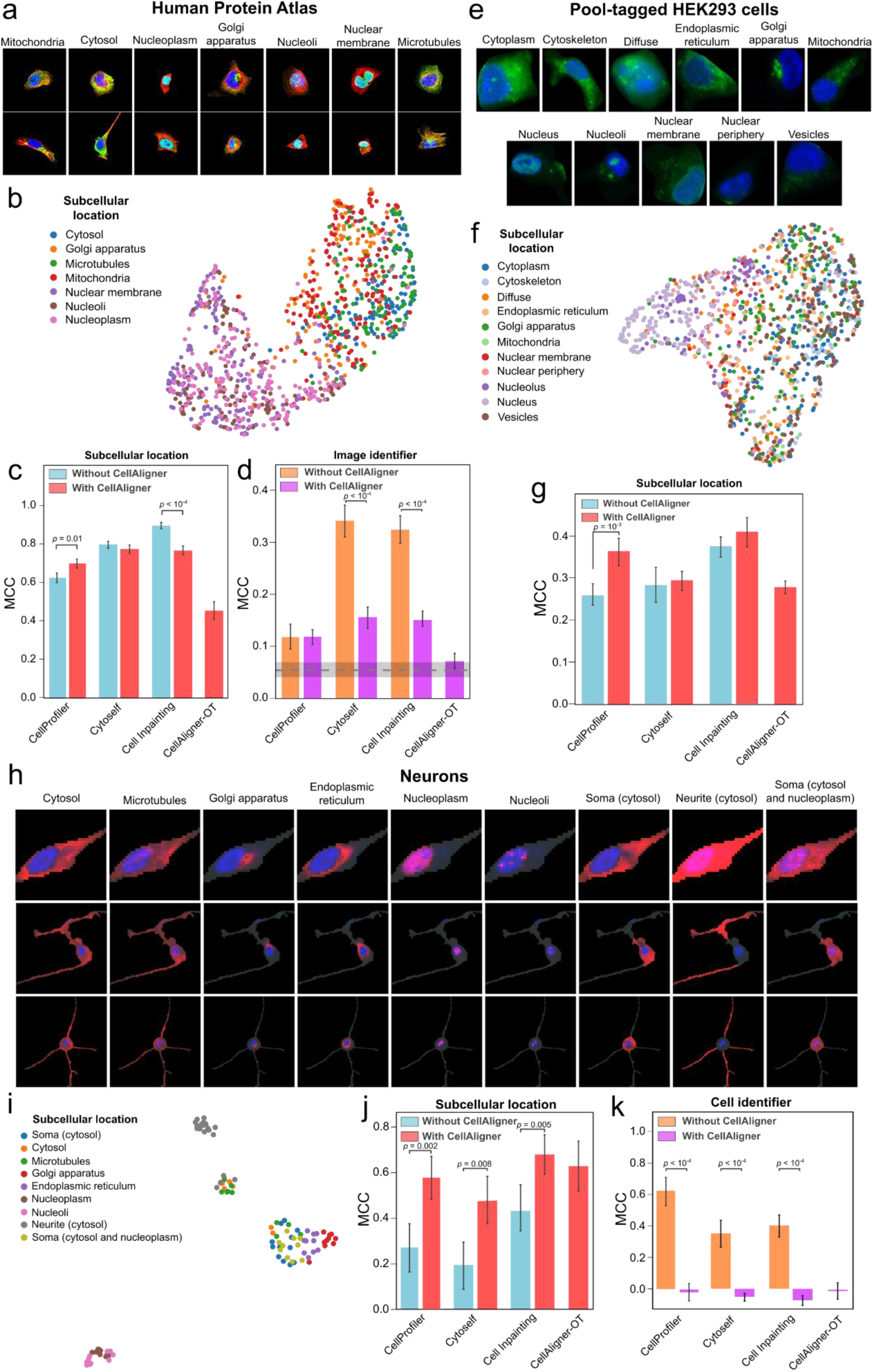
Quantitative evaluation of CellAligner. **(a-d)** Evaluation of CellAligner using images from the Human Protein Atlas. (**a**) Representative test images showing seven manually annotated subcellular localization patterns. (**b**) UMAP embedding of the subcellular localization space produced by CellAligner-OT, colored by ground-truth localization labels. (**c**) MCC of k-nearest neighbor classifiers trained to predict subcellular localization labels from the outputs of CellAligner-OT and three localization analysis methods applied either to the original images or to the anchor-space representations generated by CellAligner. (**d**) MCC of k-nearest neighbor classifiers trained to predict image identity from the same outputs. The dashed line and grey band indicate the baseline performance, defined by the average MCC ± standard deviation of a random classifier. This baseline MCC is greater than zero because images contain multiple cells with the same localization pattern, allowing partial inference of image identity from localization alone. **(e-g)** Evaluation using images from pool-tagged HEK293 cells. (**e**) Representative test images depicting eleven annotated subcellular localization patterns. (**f**) UMAP embedding of the subcellular localization space generated by CellAligner-OT, colored by ground-truth labels. (**g**) MCC of k-nearest neighbor classifiers trained to predict subcellular localization labels from the outputs of CellAligner-OT and three localization analysis methods applied either to the original images or to the anchor-space representations generated by CellAligner. **(h-l)** Evaluation using images from neurons with simulated subcellular localization patterns. (**h**) Representative test images showing nine subcellular localization patterns. (**i**) UMAP embedding of the subcellular localization space generated by CellAligner-OT, colored by ground-truth localization labels. (**j**) MCC of k-nearest neighbor classifiers trained to predict subcellular localization labels from the outputs of CellAligner-OT and three localization analysis methods applied either to the original images or to the anchor-space representations generated by CellAligner. (**k**) MCC of k-nearest neighbor classifiers trained to predict cell identity from the same outputs. Two-sided Wilcoxon rank-sum test p-values are shown when statistically significant (𝑝 ≤ 0.05).

In this analysis, Cytoself and Paired Cell Inpainting achieved the highest MCCs in predicting subcellular localization labels, significantly outperforming the other methods (**Fig. 2c**). This strong performance is expected in this setting, given both the scale of their training datasets and their close similarity to the test data. Remarkably, Paired Cell Inpainting maintained high performance even when initialized with random weights and left untrained, suggesting that part of its performance originates from architectural priors intrinsic to convolutional networks rather than from learned representations alone (**Supplementary Fig. 3a**). Similar high baselines for convolutional networks have been documented in other contexts^29,30^, raising the possibility that Paired Cell Inpainting, and potentially Cytoself, may exploit image-associated features that are not specific to subcellular protein localization.

Consistent with this possibility, both Cytoself and Paired Cell Inpainting achieved high MCCs for predicting cell-line identity and image identity from the subcellular localization space, even after conditioning on subcellular localization label (**Fig. 2d** and **Supplementary Fig. 3b**, adjusted MCC = 0.56 and 0.52 for cell line, and 0.29 and 0.27 for image ID; permutation test *p*-value < 10^-4^ in all cases). CellProfiler also performed above baseline for predicting cell-line and image identity (adjusted MCC = 0.23 and 0.06 for cell line and image ID, respectively; permutation test *p*-value < 10^-4^), indicating that morphology, cell-line-specific properties, and other image-associated features influence the representations produced by this method.

Taken together, these results suggest that, in Human Protein Atlas data, current methods can accurately predict broad localization labels while still encoding substantial morphology- and image-associated variation.

### CellAligner reduces morphological confounding and improves subcellular localization analysis

To quantitatively assess the utility of CellAligner for reducing the confounding effects of morphological variation, we repeated the Human Protein Atlas benchmarking analysis above using the anchor-cell representations produced by CellAligner instead of the original images. In addition to CellProfiler, Cytoself, and Paired Cell Inpainting, we considered the subcellular localization space obtained by computing OT distances between mapped protein distributions within CellAligner anchor-cell representations (CellAligner-OT).

Overall, applying current methods to the anchor-cell representations produced by CellAligner substantially reduced their dependence on cell-line and image identity relative to the original images (**Fig. 2d** and **Supplementary Fig. 3b**). Similarly, the MCC of CellAligner-OT for predicting cell-line and image identity was close to the baseline (adjusted MCC = 0.09 and 0.01 for cell line and image ID, respectively; permutation test *p*-value < 10^-4^ and 0.06).

Despite these reductions in cell-line- and image-associated confounding, localization-label prediction was not substantially affected by replacing the original images with CellAligner anchor-cell representations (**Fig. 2b, c**), except for Paired Cell Inpainting, whose MCC was slightly reduced after applying CellAligner, potentially due to overfitting, since the model was originally trained using the Human Protein Atlas dataset.

Because localization labels are not equally represented across cell lines in the Human Protein Atlas, subcellular localization and morphological variation are not fully independent in this dataset. To better isolate the effect of morphology-associated confounding on subcellular protein localization analysis, we next considered a recently published pooled protein-tagging dataset of HEK293 cells^31^, consisting of 1,150 cells and 46 tagged proteins (**Fig. 2e**). As above, we assessed the ability of each method to distinguish the manually annotated localization patterns for each tagged protein using either the original cell images or the anchor-cell representations produced by CellAligner (**Fig. 2f** and **Supplementary Fig. 2**). In this analysis, the MCC of all current methods increased when applied to CellAligner anchor-cell representations, with particularly marked improvements for CellProfiler (**Fig. 2g**).

Reasoning that morphological confounding should be especially severe in cells with complex morphologies, such as neurons, astrocytes, or macrophages, we next generated a semi-synthetic dataset of neuronal cell images exhibiting various subcellular localization patterns that were independent of neuronal morphology (**Fig. 2h**). This dataset was constructed by mapping organelle-specific protein localization patterns from the Human Protein Atlas onto imaged neuronal morphologies (Methods). As in the previous analyses, we quantified the ability of each method to discriminate subcellular localization patterns independently of neuronal morphology.

In this dataset, all current methods performed poorly on the original neuronal images, with low MCCs for localization-label prediction and strong encoding of cell morphology, consistent with the challenges posed by neuronal morphologies in subcellular localization analysis (**Fig. 2j**). However, when applied to the anchor-cell representations produced by CellAligner, localization-label prediction improved substantially for all methods, with an average 2.1-fold increase in MCC relative to their performance on the original images (**Fig. 2j**). This performance was comparable to that achieved by CellAligner-OT (**Fig. 2i, j**). At the same time, morphology-associated confounding was reduced to baseline levels (**Fig. 2k** and **Supplementary Fig. 4**).

Together, these results demonstrate that CellAligner separates subcellular protein localization from morphological variation in cells with complex architectures, enabling morphology-robust analysis of subcellular localization and substantially improving accuracy of existing methods in challenging settings such as neurons.

### CellAligner detects alterations in neuronal lysosomal transport

To assess the ability of CellAligner to identify differences in subcellular localization under complex, biologically relevant conditions, we tagged the lysosomal marker lysosome-associated membrane protein 1 (LAMP1) in 2-week-old human iPSC-derived neurons treated with either a lysosomal cholesterol export inhibitor (U18666A, 5 μg/mL) or DMSO (**Fig. 3a**). Lysosomes are critical in cholesterol metabolism, and alterations in cholesterol transport are implicated in multiple neurodegenerative diseases^32^, such as Niemann-Pick disease type C (NPC)^33^. U18666A inhibits the cholesterol transporter activity of the NPC1 protein, leading to the accumulation of unesterified cholesterol in late lysosomes and impaired microtubule-based transport^34^. In neurons, this perturbation results in defective lysosomal trafficking along neurites, a phenotype that is easily masked by the considerable morphological variability and structural complexity of neurons.

**Figure 3.**
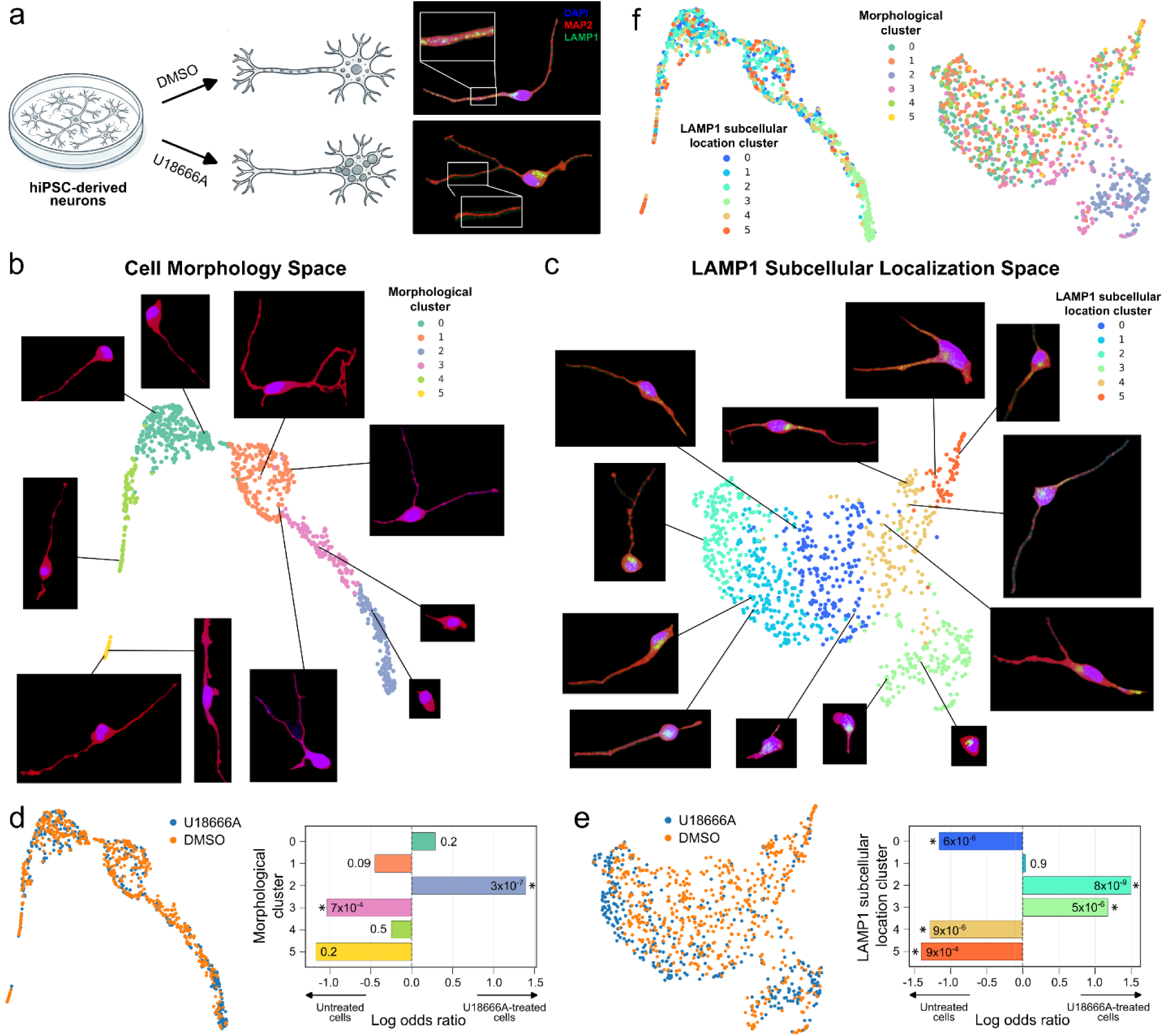
Zero-shot identification of impaired cholesterol transport in human iPSC-derived neurons using CellAligner-OT. (**a**) Schematic of the experimental design. Human iPSC-derived neurons are treated two weeks post-induction with DMSO or the lysosomal cholesterol export inhibitor U18666A and imaged for DAPI, MAP2, and LAMP1. This treatment disrupts lysosomal trafficking, marked by LAMP1, along the neurites. (**b**) UMAP representation of the cell morphology space, colored by morphological clusters. Representative cell images are shown. (**c**) UMAP representation of the LAMP1 subcellular localization space, colored by localization clusters. Representative cell images are shown. (**d**) UMAP of the cell morphology space colored by treatment (left) and results of Fisher’s exact test enrichment analysis in each morphological cluster (right). Benjamini-Hochberg-adjusted *p*-values are indicated. (**e**) UMAP of the LAMP1 subcellular localization space colored by treatment (left) and results of Fisher’s exact test enrichment analysis in each localization cluster (right). Benjamini-Hochberg-adjusted *p*-values are indicated. (**f**) UMAPs of the cell morphology (left) and LAMP1 subcellular localization (right) spaces, colored by morphological and subcellular localization clusters, respectively.

We imaged and segmented 948 neurons and applied CellAligner-OT to compute morphology and LAMP1 subcellular localization summary spaces. The morphology space segregated neurons according to structural features, with cells differing in neurite number occupying distinct regions (**Fig. 3b**). In contrast, the LAMP1 localization space separated neurons exhibiting soma-restricted lysosomal localization from those with lysosomes extending into neurites (**Fig. 3c**).

To quantify treatment-induced changes in neuronal morphology and lysosome localization, we clustered each space and tested for enrichment of control and U18666A-treated neurons in each cluster (**Fig. 3d-f**). Consistent with impaired lysosome trafficking, CellAligner-OT identified a significant overrepresentation of U18666A-treated neurons in clusters characterized by soma-restricted lysosomal localization (odds ratio = 2.3 - 2.8, Fisher exact test *q*-value < 10^-5^). Additionally, CellAligner’s morphology space identified a small subset of neurons with rounded morphology significantly associated with U18666A treatment (odds ratio = 2.6, Fisher exact test *q*-value = 7 x 10^-4^), likely reflecting treatment-induced cellular stress or death.

Together, these results demonstrate that CellAligner can be used to robustly resolve and attribute biologically meaningful changes in both complex protein localization patterns and associated neuronal morphology, even in systems with substantial intrinsic variability.

### Deep metric learning scales CellAligner to label-free datasets consisting of millions of cells

CellAligner involves solving an expensive computational optimization problem, which limits its scalability to large imaging datasets. Computing an anchor-cell representation for the 1,021 cells in our benchmarking Human Protein Atlas dataset required 1.4 hours on a standard 8-core desktop computer, while computing all pairwise OT distances between the mapped protein distributions took an additional 3.5 hours. However, high-throughput imaging-based methods^5-7,35,36^ can profile dozens to hundreds of proteins across hundreds of thousands of cells.

To enable CellAligner-based analysis in high-throughput settings, we developed deep CellAligner-OT (dCellAligner-OT), a deep metric learning framework that efficiently approximates CellAligner-OT subcellular protein localization distances and anchor-cell representations of protein distributions (**Fig. 4a**; Methods).

**Figure 4.**
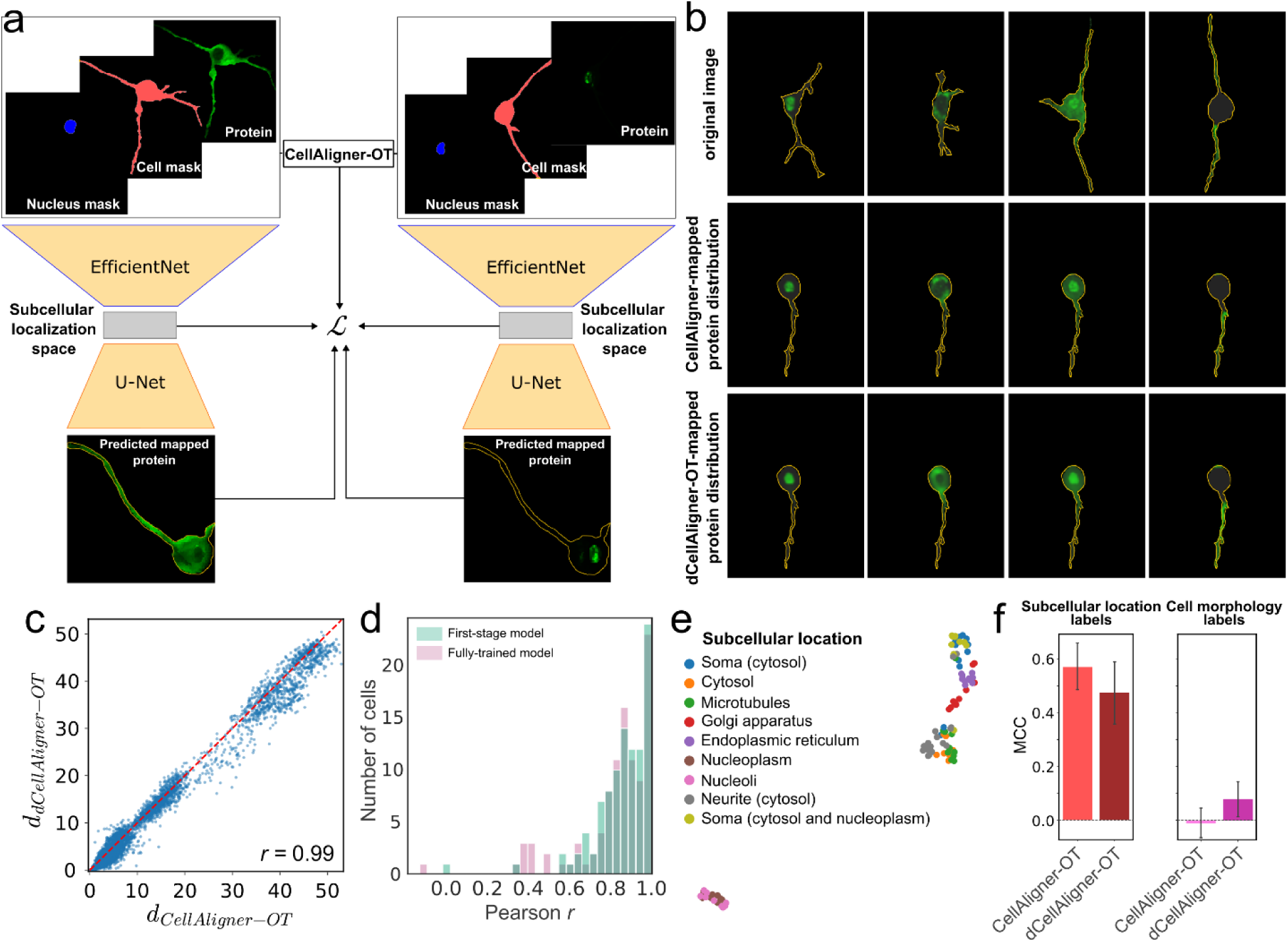
A deep metric learning model for scalable, metric geometry-based analysis of subcellular protein localization. (**a**) Schematic of the dCellAligner-OT Siamese architecture. The encoder takes as input the cellular and nuclear segmentation masks and the protein stain for each cell. The model is optimized using a metric-learning loss that penalizes discrepancies between CellAligner-OT distances computed for pairs of cells and the distances between their corresponding embeddings. This loss is regularized by a reconstruction objective implemented with a decoder, which directs the embeddings to retain information necessary to reconstruct the mapped protein distribution on an anchor cell. (**b**) Examples of neuronal protein stains mapped onto a reference anchor cell using CellAligner-OT or the trained dCellAligner-OT model. (**c**) Scatter plot comparing pairwise CellAligner-OT distances with pairwise dCellAligner-OT distances in the semi-synthetic neuronal test dataset. (**d**) Distribution of pixel-wise Pearson correlation coefficients between CellAligner-OT and dCellAligner-OT mapped protein distributions, for both first-stage and fully-trained dCellAligner-OT models. (**e**) UMAP representation of the subcellular localization space produced by CellAligner-OT for the semi-synthetic neuronal test dataset, colored by ground-truth subcellular localization labels. (**f**) MCC of k-nearest neighbor classifiers trained to predict subcellular localization (left) and cell morphology (right) labels based on the subcellular localization space produced by dCellAligner-OT for the semi-synthetic neuronal test dataset.

The dCellAligner-OT architecture consists of a convolutional EfficientNet encoder^37^ that takes as input the cellular and nuclear segmentation masks and the protein-staining image for each cell, together with a U-Net decoder^38^ that produces an anchor-cell representation of the protein distribution that can be analyzed using existing cell image analysis methods. The model is trained using a two-stage procedure (Methods). In the first stage, the model is optimized using a reconstruction objective that encourages the embeddings to retain the information necessary to reconstruct the mapped protein distribution in the anchor cell. In the second stage, the model is fine-tuned as a Siamese network using an additional metric-learning loss that penalizes discrepancies between CellAligner-OT distances computed for pairs of cells and Euclidean distances between their corresponding embeddings. Thus, in the fully trained model, Euclidean distances in the low-dimensional dCellAligner-OT embedding approximate the corresponding CellAligner-OT distances between pairs of cells. In addition, the metric-learning stage regularizes the anchor-cell reconstructions, improving the quality of the mapped representations.

We evaluated dCellAligner-OT using the Human Protein Atlas and semi-synthetic neuronal benchmarking datasets. To this end, we trained separate instances of the model on 15,487 cells from the Human Protein Atlas and 15,331 semi-synthetic neurons, with no overlap between training and test data. We assessed model performance by using the resulting embedding spaces to predict ground-truth localization labels with a k-nearest neighbor classifier. In both datasets, dCellAligner-OT closely approximated CellAligner-OT distances between cells (Pearson *r* = 0.99, two-sided t-test *p*-value < 10^−16^) and accurately reconstructed the corresponding CellAligner-mapped protein distributions (**Figs. 4b-d** and **Supplementary Figs. 5a-c**).

In the Human Protein Atlas dataset, the median pixel-wise Pearson correlation between CellAligner and dCellAligner-OT anchor-cell reconstructions was *r* = 0.81 for the first-stage model and *r* = 0.90 for the fully trained model (two-side Wilcoxon rank-sum test *p*-value = 10^-83^). Reconstruction correlations were statistically significant for 99.7% and 99.8% of the cells, respectively (permutation test *p*-value < 0.05; **Supplementary Figs. 5c**). In this dataset, dCellAligner-OT achieved MCC values comparable to CellAligner-OT for distinguishing localization patterns (MCC = 0.52 vs 0.45), while exhibiting reduced sensitivity to image features unrelated to subcellular localization (**Supplementary Fig. 5d, e**).

In the neuronal benchmarking dataset, the median pixel-wise Pearson correlation between CellAligner and dCellAligner-OT anchor-cell reconstructions was *r* = 0.86 for both the first-stage and fully trained models (**Fig. 4b**). In this dataset, dCellAligner-OT yielded slightly lower MCC values than CellAligner-OT (MCC = 0.47 vs 0.57), but nevertheless outperformed all existing methods while showing limited encoding of morphological variation (**Figs. 4e, f**).

Notably, dCellAligner-OT provides a substantial computational advantage. Once trained, the model computes dCellAligner-OT embeddings for 100,000 cells in approximately 9 minutes, compared to an estimated 31 days required for exhaustive pairwise CellAligner-OT computation. Moreover, dCellAligner-OT learns directly from CellAligner-OT, not requiring any labels or additional training data. Together, these results demonstrate that dCellAligner-OT enables scalable, metric geometry-based analysis of subcellular protein localization without substantially compromising accuracy, opening its application to atlas-scale studies of spatial proteomics.

## Discussion

The spatial organization of proteins within cells is a fundamental layer of cellular regulation. However, quantitatively comparing subcellular localization across morphologically complex and heterogeneous cells, such as differentiated and primary cells, remains challenging. Our analyses indicate that widely used methods for subcellular localization analysis, including CellProfiler^10^, Cytoself^14^, and Paired Cell Inpainting^13^, encode morphology- and image-associated variation that is not specific to protein localization. While this confounding has limited impact on broad localization-label prediction in relatively simple cell lines, it severely reduces performance in cells with complex morphologies. To address this limitation, we developed CellAligner, a compartment-aware cell registration framework based on fused unbalanced Gromov-Wasserstein couplings that maps protein distributions from morphologically distinct cells into shared anchor-cell morphologies while allowing partial matching. We implemented CellAligner as part of CAJAL^21^, our open-source library for metric-geometry-based analysis of cell morphology and organization.

Several observations emerge from our benchmarking analyses. First, self-supervised deep learning methods such as Cytoself and Paired Cell Inpainting achieve strong performance on morphologically simple cell lines, such as those in the Human Protein Atlas^5^ and OpenCell-like^6^ datasets, but their embeddings also encode substantial cell-line- and image-associated variation. Removing this variation with CellAligner does not necessarily improve localization-label prediction in these datasets, likely because these methods were developed and trained in closely related imaging contexts. Second, applying CellProfiler to CellAligner anchor-cell representations rather than to the original images substantially reduces its dependence on cellular morphology and improves its ability to distinguish subcellular localization patterns across all datasets we tested. This suggests that classical feature-based methods can become competitive with more complex representation-learning approaches when morphology-associated variation is first removed. Third, in morphologically complex cells, all evaluated current methods perform poorly when applied directly to the original images, with representations strongly confounded by cell shape. In this setting, applying these methods to CellAligner anchor-cell representations substantially improved localization-label prediction and reduced morphology-associated confounding to near-baseline levels. In these analyses, Paired Cell Inpainting applied to CellAligner representations and CellAligner-OT were the top-performing approaches, closely followed by CellProfiler applied to CellAligner representations. These results indicate that CellAligner can extend existing image-analysis methods to challenging cellular contexts without requiring the development of specialized intracellular coordinate systems^15,16^, which may be difficult or impossible to define for highly complex and heterogeneous cell morphologies. We illustrated these capabilities by analyzing lysosomal trafficking alterations in iPSC-derived neurons after treatment with U18666A, a cholesterol transport inhibitor^34^.

To scale CellAligner to datasets containing millions of cells, we further developed dCellAligner-OT, a deep metric learning model trained to approximate the localization distances and anchor-cell representations produced by CellAligner-OT. This distillation strategy provided up to a 5,000-fold speedup relative to exact CellAligner-OT while largely preserving accuracy. Related teacher-student strategies have previously been used to approximate Wasserstein or Gromov-Wasserstein distances in point-cloud settings^39,40^. Here, we extend this idea to a more complex biomedical imaging problem in which the inputs include multi-channel microscopy images and segmentation masks, partial matching is required, and the output consists of protein distributions represented within a shared anchor-cell morphology. More broadly, our application demonstrates the potential of algorithmic distillation to make computationally intensive geometric methods practical for large-scale biomedical imaging datasets.

Our current implementation of CellAligner has several limitations. First, exact GW computations scale poorly with the number of pixels, which often requires anchor-cell representations to be computed at lower resolution than the original images. This may limit sensitivity to localization patterns that differ only at very fine spatial scales. Second, for the same reasons, the computation of GW correspondences becomes computationally challenging in 3D imaging data, and CellAligner is currently restricted to 2D microscopy images. We hope to address both limitations in future work.

Overall, CellAligner provides a broadly applicable framework for morphology-robust quantification of subcellular protein organization. By separating localization differences from variation in cell shape, CellAligner enables quantitative analysis of intracellular organization in morphologically complex cells and provides a morphology-robust scalable approach for image-based studies of disease mechanisms, drug responses, and biomarker discovery.

### Methods CellAligner

### method

CellAligner takes as input cell segmentation masks, including morphological stain intensities (i.e., nuclear or membrane stains), alongside images of one or more fluorescently labelled proteins. Given a cell, 𝑖, CellAligner first solves a fused unbalanced GW problem to map the interior of the cells to an anchor cell 𝑗 based on the segmentation masks and morphological stain intensities^18^

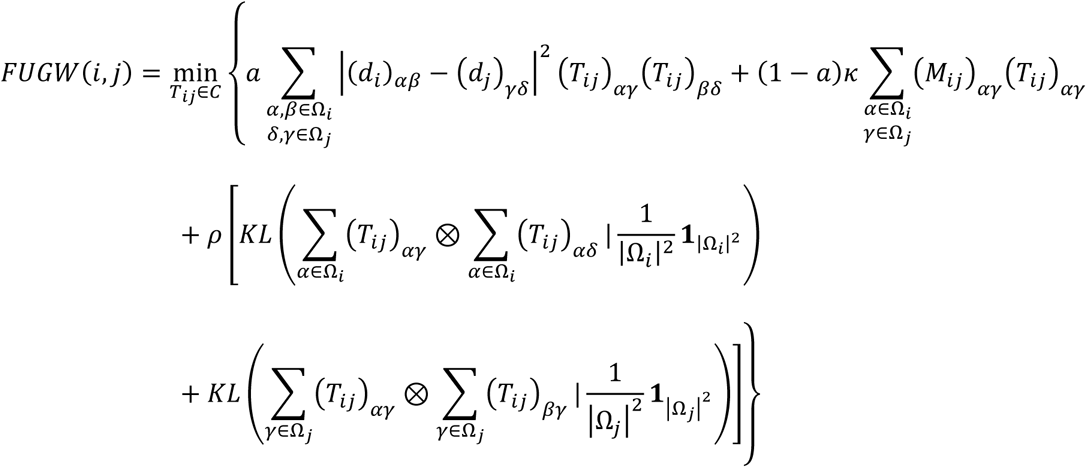

where the matrix 𝑇_𝑖𝑗_ specifies a weighted correspondence between the pixels in the segmentations masks Ω_𝑖_ and Ω_𝑗_, 𝐶 denotes the space of all possible weighted correspondences, (𝑑_𝑖_)_𝛼𝛽_ is the geodesic distance between pixels 𝛼 and 𝛽 within Ω_𝑖_, (𝑀_𝑖𝑗_)_𝛼𝛾_ is the dissimilarity in morphological stain intensities between pixels 𝛼 ∈ Ω_𝑖_ and 𝛾 ∈ Ω_𝑗_, 𝜅 is a user-specified normalization parameter, 𝑎 ∈ [0,1] and 𝜌 ∈ [0, ∞) are user-specified parameters controlling the relative contribution of morphological staining information to the correspondence and the penalty of leaving parts of the cell unmapped, respectively, ⊗ is the Kronecker product between vectors, 𝟏_𝑁_ is a vector of dimension 𝑁 with all components equal to one, and 𝐾𝐿(⋅ | ⋅) denotes the Kullback-Leibler divergence. By incorporating nuclear segmentation information, the resulting maps are constrained to preserve subcellular compartmentalization (e.g., avoiding the mixing of cytoplasmic and nuclear compartments). This helps prevent the emergence of multiple nearly degenerate mappings for highly symmetric shapes (e.g., rounded cells). Because the distance matrices 𝑑_𝑖_ are inherently invariant to rotations and translations, the computation of GW correspondences does not require pre-aligning the cells. We use the Python Optimal Transport library^41^ to solve the fused unbalanced GW problem using the conditional gradient algorithm. For datasets with simple cell morphology, such as those from the Human Protein Atlas, it is convenient to set 𝜌 = ∞ and solve the corresponding fused GW problem.

The correspondences 𝑇_𝑖𝑗_ obtained from the above procedure are then used to map the protein distributions across cells. Specifically, given the normalized pixel intensities of a fluorescently labeled protein in cell 𝑖, (𝑔_𝑖_)_𝛼_, the mapped protein distribution in the anchor cell 𝑗 is given by

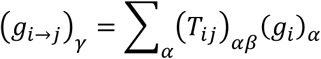

The discrepancy between the protein localization patterns of two cells 𝑖_1_ and 𝑖_2_ is computed using either a colocalization metric or a distance between feature vectors produced by a cell image analysis method. By default (CellAligner-OT), we use the Wasserstein distance between the mapped protein distributions in the anchor cell 𝑗,

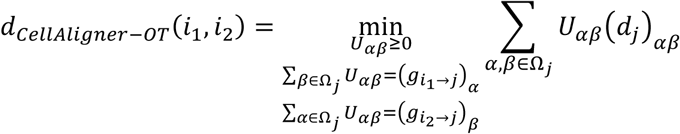

The resulting distance 𝑑_𝐶𝑒𝑙𝑙𝐴𝑙𝑖𝑔𝑛𝑒𝑟−𝑂𝑇_ is thus sensitive only to changes in the relative spatial distribution of the protein within the cell and is invariant to morphological differences between the original cells.

### Compartment-specific mapping

By utilizing morphological staining information to find optimal probabilistic cell-to-cell mappings, CellAligner tries to align the cellular components defined by the morphological stains. By default, CellAligner probabilistically weighs each cell’s pixels equally, which preserves the relative size of cellular compartments after mapping. However, in cases where the relative size of cellular compartments (cytoplasm, nucleus, etc.) is drastically different between cells, this can result in poor alignment of those compartments after mapping. To address this, we provide an option for CellAligner to perform compartment-specific mapping. For compartment-specific mapping, CellAligner additionally takes as input segmentation masks for 𝑃 cellular compartments. For a given cellular compartment 𝑝 ∈ 𝑃, CellAligner then enforces that pixels belonging to compartment 𝑝 in cells 𝑖 and 𝑗 receive equal total probability. Formally, if 𝑝_𝑖_ and 𝑝_𝑗_ denote the sets of pixels in compartment 𝑝 of cells 𝑖 and cell 𝑗, respectively, then

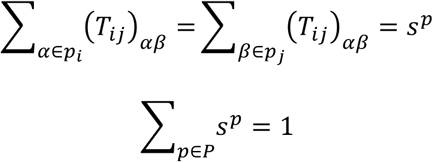

where 𝑠^𝑝^ are user-specified parameters representing the probabilistic mass assigned to each compartment. This reweighting allows CellAligner to more accurately align cellular compartments during morphological mapping. For all applications using compartment-specific weighting in this paper, we consider two cellular compartments: the cytoplasm and the nucleus.

### Integration of multiple anchor cells and proteins

We adapted the weighted nearest-neighbors algorithm from Hao *et al.*^28^ to integrate multiple subcellular localization spaces. For this purpose, we utilize the Manifold-Adaptive Dimension Estimation (MADA) algorithm^42^ to estimate the intrinsic dimensionality of each space from its pairwise distance matrix. Each distance matrix is then embedded into a Euclidean space of the estimated dimension using Isomap^43^. Finally, the resulting Euclidean spaces are integrated using the weighted-nearest neighbors algorithm. This procedure has been implemented as a function in our Python library CAJAL^21^.

### Morphology space

While the value of the *FUGW* cost function at its minimum defines a morphological distance, solving the fused unbalanced GW problem for every pair of cells becomes computationally too expensive for datasets containing more than a few hundred cells. Therefore, to quantify the variation in cell morphology, we instead followed the simpler approach described in Govek *et al*.^21^, which consists of uniformly sampling points from the cell boundary and computing the GW distance between the sampled point sets for each pair of cells, as implemented in the Python package CAJAL. This approach substantially improves the runtime of morphology space computations without significantly affecting accuracy.

### Human Protein Atlas dataset

We utilized images from version 23.0 of the Human Protein Atlas. For the initial evaluation of CellAligner, we considered a dataset of 387 cells from 58 images across 58 proteins localized to 5 manually annotated compartments (cytosol, Golgi apparatus, nucleoli, nucleoplasm, plasma membrane), and 4 human cell lines (A-431, U2OS, U-251MG, A-549). We performed cell and nuclear segmentation using the pretrained U-Net model from the Python library `hpacellseg`. We then sampled 100 points from the boundary of each cell segmentation mask and used CAJAL v1.5 to construct the cell morphology space based on the Euclidean distances between the sampled points. To construct the protein localization space, we downsampled each segmented cell image to roughly 1,000 pixels. To choose a representative set of anchor cell morphologies, we clustered the cell morphology space using the Leiden algorithm and selected the centroid cell from each cluster. We additionally included the overall centroid of the morphology space as an anchor. We performed compartment-specific mapping (𝑠^𝑐𝑦𝑡𝑜^ = 𝑠^𝑛𝑢𝑐^ = 0.5) of each cell’s subcellular protein distribution onto each anchor cell morphology using fused GW (𝜌 = ∞) with parameters 𝑎 = 0.1 and 𝜅 = 10. Intracellular distances were measured using Euclidean distance, and the binarized nuclear segmentation mask was used to compute 𝑀_𝑖𝑗_. We then applied the weighted nearest-neighbors approach described in subsection “*Integration of multiple anchor cells and proteins*” with 𝑘 = 5 to construct an integrated protein subcellular localization space from the localization spaces associated with individual anchors.

To evaluate the performance of selected existing methods for subcellular protein localization analysis (CellProfiler, Cytoself, and Paired Cell Inpainting), we analyzed a subset of the Human Protein Atlas comprising 1,043 cells from 140 images across 21 proteins localized to seven subcellular compartments (cytosol, Golgi apparatus, microtubules, mitochondria, nuclear membrane, nucleoli, and nucleoplasm) across three human cell lines (A-431, U2OS, and U-251MG).

For each method, we extracted features from individual cell images and trained 5-nearest neighbor classifiers using 10-fold cross-validation to predict annotated subcellular localization pattern, cell line identity, and image ID. We quantified performance using Mathew’s correlation coefficient (MCC). To assess statistical significance, we compared the observed MCC values to null distributions generated from 10,000 random permutations of the labels. Because labels from different categories are not independent, allowing partial inference of cell line identity and image ID from subcellular localization alone, we constructed the null distributions by only permuting labels within each subcellular localization class.

#### CellProfiler

We used CellProfiler v4.2.5 to extract features related to subcellular protein localization. Cell and nuclear segmentation masks were provided as additional inputs. Image features were extracted using the following modules and parameters:

MeasureObjectIntensityDistribution (bin number = 4); MeasureTexture (gray levels = 256; texture scales = 1-8); MeasureGranularity (subsampling factor for granularity and background = 0.25; radius of structuring element = 10; radius of granular spectrum = 16); and MeasureColocalization (maximum threshold intensity = 15), yielding a total of 514 features. We z-scored features before classification, removing those exhibiting no variance or high correlation with other features (Pearson correlation coefficient > 0.9).

#### Cytoself

We used the Cytoself model (TensorFlow implementation) trained on protein and nuclear images from the full OpenCell dataset to construct embeddings of protein localization features. Input protein and nuclear stain images were resized to 100 x 100 pixels. For experiments involving an untrained Cytoself model, we used the newer PyTorch implementation, initialized with the following parameters: input shape = 2 x 100 x 100; embedding shapes = (25, 25) and (4, 4); embedding dimension = 64; number of embeddings = 2,048; fully connected (FC) output index = all; number of classes = 100; and FC input type = vqvec. The encoder output was used for downstream analysis.

#### Paired Cell Inpainting

We used the Paired Cell Inpainting model trained on the full Human Protein Atlas to construct embeddings of protein localization features. Protein, nuclear, and microtubule stain input images were resized to 64 x 64 pixels. For datasets without microtubule imaging (HeLa pooled protein tagging, iPSC-derived neuron lysosome tagging, and semi-synthetic neuron images) we replaced the microtubule image with the cell segmentation mask.

For experiments using an untrained Paired Cell Inpainting model, we initialized the model with the following parameters: input X shape = (64, 64, 3); input Y shape = (64, 64, 2); batch size = 64; optimizer = Adam; learning rate = 1 × 10^−4^; 𝛽_1_ = 0.5; and target layer = conv3_1. For fine-tuning experiments, the Paired Cell Inpainting model was trained on 700 cell images from the HeLa pooled protein tagging dataset for 5, 10, 20, 30, and 100 epochs. The encoder output was used for downstream analysis.

### HEK293 pooled protein tagging

We downloaded a subset of cell images, corresponding cell and nucleus segmentation masks, and associated metadata from Sansbury et al.^31^, comprising 1,150 cells images across 46 tagged proteins localized to eleven annotated subcellular compartments (cytoplasm, cytoskeleton, diffuse, endoplasmic reticulum, Golgi apparatus, mitochondria, nucleus, nucleoli, nuclear membrane, nuclear periphery, and vesicles). We followed the same procedure used for the Human Protein Atlas dataset for CellProfiler, Cytoself, and Paired Cell Inpainting.

### iPSC-derived neuron lysosome tagging

WTC11 human iPSC culture protocols and neuronal differentiation protocols were performed as previously described^44^. WTC11 iPSCs (engineered with the inducible Ngn2 from the AAVS1 safe harbor locus) were maintained in mTesR1+ media (StemCell Technologies) on hESC Matrigel (Corning, #354277) coated plastic plates. iPSCs were differentiated to glutamatergic neurons as previously described^45^. Briefly, iPSCs were singularized using StemPro Accutase (5-10 min, 37C), neutralized with mTesR1+ media, and then gently pelleted at 300*g* for 5 minutes. iPSCs were resuspended to 0.7x10^6^ cells/mL in neuronal induction media containing: KnockOut DMEM/F-12 (Gibco# 12660012) + 1x NEAA (Gibco#11140050) + 1x GlutaMAX (Gibco# 35050-061) + 10 nM Y27632 ROCK inhibitor (Tocris# 125410) + 1x N2 max (Gibco #17502-048) + 2 µg/mL doxycycline hydrochloride (Sigma# D9891). This same media was replenished the following day, and again the next day except without Y27632 ROCK inhibitor. At 3 days post-induction, neural progenitors (NPCs) were dissociated with pre-warmed StemPro Accutase (10 min, 37C), gently pelleted for 5 min at 300*g*, and re-plated as indicated on glass-bottom plates. These plates were first treated with 1N HCl for 15 min, washed 5 times, coated overnight at 37C with poly-L-ornithine hydrobromide (0.1 mg/mL in borate buffer pH 8.5; Sigma, #P3655), then washed an additional 3 times before drying overnight at room temperature. iNeurons were replated in media containing: Neurobasal-A (Gibco# 10888-022) + 1x NEAA + 1xGlutaMAX + 10 nM Y27632 ROCK inhibitor + 2 µg/mL doxycycline hydrochloride + 1x B-27 plus (Gibco# A3582801) + CultureOne Supplement (Gibco #A3320201) + 10 ng/mL BDNF (PeproTech #450-02) + 10 ng/mL NT-3 (PeproTech #450-03) + 1 µg/mL laminin (Gibco# 23017-015) + 200 µM L-ascorbic acid (Sigma #A8960).

WTC11-NGN2 iPSCs were transduced with FUW-LAMP1-mCherry2 and FUW-EGFP, then sorted using a BD FACS ARIA Fusion into a polyclonal population that was positive for both mCherry2 and EGFP. These iPSCs were differentiated to iNeurons as described above. 3 days post-induction, neural progenitors (NPCs) were lifted and mixed with wildtype WTC11 (not expressing LAMP1-mCherry2 or EGFP) at a 1:20 ratio, yielding a 5% sparsely labeled population of NPCs expressing both LAMP1-mCherry2 and EGFP. This sparsely labeled population was sub-plated on a PLO-coated glass 6-well plate at 100,000 cells/well in complete neuronal maturation media. At 8 days post-induction, neurons were treated with either DMSO or 10µg/mL U18666A (Sigma, #662015) for 24 hours, then fixed (at *DPI 9*) in 2% paraformaldehyde for 15 minutes at room temperature, followed by microscopy (3 Z-stacks in 0.5µm increments) to tile each well using a Nikon Eclipse Ti2 spinning disc confocal microscope.

To perform cell and nuclear segmentation, we independently binarized the MAP2 and DAPI channels using Otsu thresholding and labeled connected components using scikit-image. To focus on the most informative morphological characteristics, we extracted an 84 µm crop centered on each cell’s soma. We removed cells with missing or multiple nuclei, as well as those in close proximity to image tile borders or neighboring neurons. After filtering, a dataset of 948 segmented neuron images was retained for downstream analysis.

For CellAligner-OT analysis of the cell images, we followed the same procedure used for the Human Protein Atlas dataset, with the following differences. The compartment-specific mapping of each cell’s subcellular protein distribution to anchor cell morphologies was performed using fused unbalanced GW with parameters 𝑎 = 0.1, 𝜅 = 10, and 𝜌 = 70 and geodesic distance, rather than fused GW. In addition, nuclear segmentation masks were incorporated to inform the mapping.

### Simulated neuron images

We generated a semi-synthetic protein imaging dataset of neurons by mapping known organelle-specific localization patterns onto distinct neuronal morphologies. Organelle-specific protein localization patterns were obtained from cell images in the Human Protein Atlas, while neuronal morphologies were derived from the iPSC-derived neuron lysosome tagging dataset.

We considered a total of 10 localization pattern-specific cell images from the Human Protein Atlas and 10 neuron cell morphologies from the lysosome tagging dataset, yielding a dataset of 100 semi-synthetic neuron images generated by mapping each localization pattern to each neuronal morphology. For most organelle patterns, mappings were performed using fused GW. For endoplasmic reticulum- and Golgi apparatus-specific patterns, fused unbalanced GW was used to produce a more realistic protein distribution around the soma.

Soma- and projection-specific location patterns were generated synthetically based on manual segmentation of the neuronal soma for each morphology, by zeroing protein fluorescence values in non-soma or soma pixels, respectively.

We used the same procedure as described for the iPSC-derived neuron lysosome tagging dataset to perform the CellAligner analysis of semi-synthetic cell images.

### Implementation and evaluation of dCellAligner-OT

#### Model architecture

The dCellAligner-OT model consists of an EfficientNet encoder 𝐸 and a U-Net decoder 𝐹. The encoder takes as input a cell image 𝑥 ∈ ℝ^𝑊×𝐻×3^ composed of three channels: the fluorescent protein distribution, the binary cell segmentation mask, and the binary nucleus segmentation mask. As with the CellAligner-OT algorithm, the input fluorescent protein channel is normalized to represent a probability distribution. The encoder utilizes the EfficientNet-B0 architecture with weights pretrained on ImageNet and maps each cell image 𝑥 to a low-dimensional latent representation 𝑧 ∈ ℝ^𝑙^. The decoder 𝐹 takes the latent representation 𝑧 as input to reconstruct the spatial distribution of protein, after mapping to the anchor cell morphology, 𝑥̂ ∈ ℝ^𝑊×𝐻×1^. To learn the CellAligner-OT metric space, the dCellAligner-OT model is organized as a Siamese network consisting of two identical networks sharing weights for 𝐸 and 𝐹, trained on pairs of cell images.

#### Loss functions

During training the dCellAligner-OT model is optimized with respect to a composite loss function ℒ_𝑡𝑜𝑡𝑎𝑙_with three terms: a distance loss (ℒ_𝑑𝑖𝑠𝑡_), a reconstruction loss (ℒ_𝑟𝑒𝑐𝑜𝑛_), and a sparsity loss (ℒ_𝑠𝑝_)

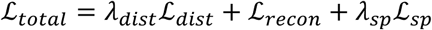

The distance loss is the Mean Squared Error (MSE) between the Euclidean distance between pairs of cell images in the latent embedding space and the true CellAligner-OT distance. This ensures the true GW-OT distances are preserved in the learned dCellAligner-OT embedding space.

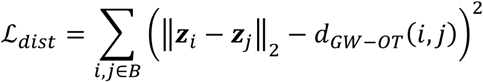

where 𝐵 is a minibatch of cells and 𝒛_𝑖_ the latent embedding of the 𝑖-th cell in the minibatch. To ensure the latent representation retains biologically relevant information spatial protein localization pattern, we include the reconstruction loss. This loss is defined as the Kullback-Leibler (KL) divergence between the true input protein distribution, after mapping to the anchor cell morphology, 𝑥^𝑚^, and the predicted mapped protein distribution 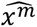, for a given cell image:

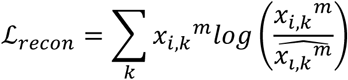

where 𝑘 indexes the pixels in the flattened protein image vector. Additionally, we regularize the model by introducing a sparsity constraint loss, ℒ_𝑠𝑝_, which imposes a penalty based on the deviates of the mean activations of the latent layer from a specified target sparsity 𝜂.

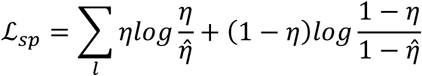

Where 𝜂̂ represents the mean activation across nodes in the latent layer.

#### Training procedure

Training is performed in a two-step process. First, the autoencoder structure (singular encoder 𝐸 and decoder 𝐹) is pre-trained on individual cell images, only optimizing the reconstruction loss. Second, the full Siamese network is trained on pairs of cell images, optimizing the full composite loss function.

#### Training dCellAligner-OT on Human Protein Atlas images

We trained the dCellAligner-OT model on 15,480 cell images from the Human Protein Atlas. To encourage the learned localization embedding space to be invariant to rigid transformations, we rotationally aligned each cell image according to the major and minor axes of the cell segmentation mask. We augmented the training images by randomly squashing and stretching each cell image along its minor axis. Each cell image was then rescaled to 64 x 64 pixels.

We trained dCellAligner-OT using the following parameters: batch size = 32; optimizer = Adam; learning rate = 1 × 10^−3^; learning-rate decay = 0.95; embedding size = 50; 𝜆_𝑑𝑖𝑠𝑡_ = 0.1; and 𝜆_𝑠𝑝_ = 1 × 10^−3^; and 𝜂 = 0.1. For the first training stage, we trained the model on the 15,480 individual cell images for 50 epochs. For the second training stage, we trained the model on 150,000 pairs of cell images for 25 epochs.

Due to the computational cost of computing GW-based mappings, we mapped each cell to a single anchor-cell morphology selected from the anchor morphologies used to benchmark CellAligner-OT on the Human Protein Atlas dataset. Additional robustness to anchor-cell could be obtained by training multiple dCellAligner-OT models using different anchor-cell morphologies and integrating the resulting subcellular localization embedding spaces using the same weighted nearest-neighbors approach used in CellAligner-OT.

#### Training dCellAligner-OT on simulated neuron images

We generated a large semi-synthetic dataset of neuronal protein images by mapping protein localization patterns from 11,725 Human Protein Atlas cell images onto 983 unique neuronal morphologies obtained from the iPSC-derived neuron lysosome-tagging dataset, as described in the subsection *“Simulated neuron images”*.

Since the CellAligner-OT analysis of the simulated neuronal protein images allowed partial mappings, we preserved the relative differences in cell size. Thus, instead of rescaling each cell image to a fixed size, as in the Human Protein Atlas dCellAligner-OT analysis, we padded each cell image to 256 x 256 pixels and did not augment the training datasets with random squashing or stretching. We centered each image on the nucleus in addition to the rotational alignment.

We trained dCellAligner-OT using the same parameters as in the Human Protein Atlas analysis. In the first training stage, we trained the model on the 11,725 individual cell images for 50 epochs. In the second training staged, we trained the model on 35,000 pairs of cell images for 25 epochs. As in the Human Protein Atlas dCellAligner-OT analysis, we used a single anchor cell.

### Code availability

The source code of CellAligner is available at https://github.com/CamaraLab/CAJAL (modules src/cajal/subcellular.py and src/cajal/subcellular_dl.py). Tutorials and documentation are available at https://cajal.readthedocs.io/.

## Data availability

All datasets used in this study are publicly available. Human Protein Atlas images are available at https://www.proteinatlas.org/humanproteome/subcellular, and the pooled-tagging HEK293 dataset is available at https://pooledtagging.org/validated-tags/. The semi-synthetic and iPSC-derived neuronal datasets generated will be deposited in Zenodo.

## Acknowledgements

This work has been supported by the U.S. National Institutes of Health (NIH) through grant RF1MH130553 from the National Institute for Mental Health (NIMH), as well as by a Collaborative Pairs Project Award (Cycle 2) from the Chan Zuckerberg Initiative.

## Author contributions

R. H. developed CellAligner and dCellAligner-OT and performed all computational analyses. N. N. N. and O. S. generated the iPSC-derived neuronal dataset. P. G. C. and R. H. conceived the project and wrote the initial draft of the manuscript, with contributions from all authors.

## References

1 Sigaeva, A., Hutchings, C., Cesnik, A., Lilley, K. S. & Lundberg, E. Subcellular localization as a driver of protein function. Nat Rev Mol Cell Biol, doi:10.1038/s41580-026-00947-3 (2026).

2 Garcia-Cazorla, A., Oyarzabal, A., Saudubray, J. M., Martinelli, D. & Dionisi-Vici, C. Genetic disorders of cellular trafficking. Trends Genet 38, 724–751, doi:10.1016/j.tig.2022.02.012 (2022).

3 Lundberg, E. & Borner, G. H. H. Spatial proteomics: a powerful discovery tool for cell biology. Nat Rev Mol Cell Biol 20, 285–302, doi:10.1038/s41580-018-0094-y (2019).

4 Lelek, M. et al. Single-molecule localization microscopy. Nat Rev Methods Primers 1, doi:10.1038/s43586-021-00038-x (2021).

5 Thul, P. J. et al. A subcellular map of the human proteome. Science 356, doi:10.1126/science.aal3321 (2017).

6 Cho, N. H. et al. OpenCell: Endogenous tagging for the cartography of human cellular organization. Science 375, eabi6983, doi:10.1126/science.abi6983 (2022).

7 Schaffer, L. V. et al. Multimodal cell maps as a foundation for structural and functional genomics. Nature, doi:10.1038/s41586-025-08878-3 (2025).

8 Roberts, B. et al. Systematic gene tagging using CRISPR/Cas9 in human stem cells to illuminate cell organization. Mol Biol Cell 28, 2854–2874, doi:10.1091/mbc.E17-03-0209 (2017).

9 Koch, B. et al. Generation and validation of homozygous fluorescent knock-in cells using CRISPR-Cas9 genome editing. Nat Protoc 13, 1465–1487, doi:10.1038/nprot.2018.042 (2018).

10 Stirling, D. R. et al. CellProfiler 4: improvements in speed, utility and usability. BMC bioinformatics 22, 433, doi:10.1186/s12859-021-04344-9 (2021).

11 Khotanzad, A. & Hong, Y. H. Invariant image recognition by Zernike moments. IEEE Transactions on pattern analysis and machine intelligence 12, 489–497 (1990).

12 Seal, S. et al. Cell Painting: a decade of discovery and innovation in cellular imaging. Nat Methods 22, 254–268, doi:10.1038/s41592-024-02528-8 (2025).

13 Lu, A. X., Kraus, O. Z., Cooper, S. & Moses, A. M. Learning unsupervised feature representations for single cell microscopy images with paired cell inpainting. PLoS computational biology 15, e1007348, doi:10.1371/journal.pcbi.1007348 (2019).

14 Kobayashi, H., Cheveralls, K. C., Leonetti, M. D. & Royer, L. A. Self-supervised deep learning encodes high-resolution features of protein subcellular localization. Nat Methods 19, 995–1003, doi:10.1038/s41592-022-01541-z (2022).

15 Le, T. et al. Cell shapes decode molecular phenotypes in image-based spatial proteomics. bioRxiv, doi:10.1101/2025.05.13.653868 (2025).

16 Viana, M. P. et al. Integrated intracellular organization and its variations in human iPS cells. Nature 613, 345–354, doi:10.1038/s41586-022-05563-7 (2023).

17 Mémoli, F. Gromov–Wasserstein distances and the metric approach to object matching. Foundations of computational mathematics 11, 417–487 (2011).

18 Thual, A. et al. Aligning individual brains with fused unbalanced Gromov Wasserstein. Advances in neural information processing systems 35, 21792–21804 (2022).

19 Vayer, T., Chapel, L., Flamary, R., Tavenard, R. & Courty, N. Fused Gromov-Wasserstein Distance for Structured Objects. Algorithms 13, 212 (2020).

20 Séjourné, T., Vialard, F.-X. & Peyré, G. The unbalanced gromov wasserstein distance: Conic formulation and relaxation. Advances in Neural Information Processing Systems 34, 8766–8779 (2021).

21 Govek, K. W. et al. CAJAL enables analysis and integration of single-cell morphological data using metric geometry. Nature communications 14, 3672, doi:10.1038/s41467-023-39424-2 (2023).

22 Fletcher, P. A., Scriven, D. R., Schulson, M. N. & Moore, E. D. Multi-image colocalization and its statistical significance. Biophys J 99, 1996–2005, doi:10.1016/j.bpj.2010.07.006 (2010).

23 Manders, E. M. M., Verbeek, F. J. & Aten, J. A. Measurement of co-localization of objects in dual-colour confocal images. J Microsc 169, 375–382, doi:10.1111/j.1365-2818.1993.tb03313.x (1993).

24 Adler, J. & Parmryd, I. Quantifying colocalization by correlation: the Pearson correlation coefficient is superior to the Mander’s overlap coefficient. Cytometry A 77, 733–742, doi:10.1002/cyto.a.20896 (2010).

25 Lagache, T. et al. Mapping molecular assemblies with fluorescence microscopy and object-based spatial statistics. Nature communications 9, 698, doi:10.1038/s41467-018-03053-x (2018).

26 Naas, J. et al. MultiMatch: geometry-informed colocalization in multi-color super-resolution microscopy. Commun Biol 7, 1139, doi:10.1038/s42003-024-06772-8 (2024).

27 Tameling, C. et al. Colocalization for super-resolution microscopy via optimal transport. Nat Comput Sci 1, 199–211, doi:10.1038/s43588-021-00050-x (2021).

28 Hao, Y. et al. Integrated analysis of multimodal single-cell data. Cell 184, 3573–3587 e3529, doi:10.1016/j.cell.2021.04.048 (2021).

29 Jarrett, K., Kavukcuoglu, K., Ranzato, M. A. & LeCun, Y. in 2009 IEEE 12th international conference on computer vision. 2146-2153 (IEEE).

30 Saxe, A. M. et al. in Icml. 6.

31 Sansbury, S. E. et al. Pooled tagging and hydrophobic targeting of endogenous proteins for unbiased mapping of unfolded protein responses. Mol Cell 85, 1868–1886 e1812, doi:10.1016/j.molcel.2025.04.002 (2025).

32 Arenas, F., Garcia-Ruiz, C. & Fernandez-Checa, J. C. Intracellular Cholesterol Trafficking and Impact in Neurodegeneration. Front Mol Neurosci 10, 382, doi:10.3389/fnmol.2017.00382 (2017).

33 Vance, J. E. & Karten, B. Niemann-Pick C disease and mobilization of lysosomal cholesterol by cyclodextrin. J Lipid Res 55, 1609–1621, doi:10.1194/jlr.R047837 (2014).

34 Lu, F. et al. Identification of NPC1 as the target of U18666A, an inhibitor of lysosomal cholesterol export and Ebola infection. Elife 4, doi:10.7554/eLife.12177 (2015).

35 Feldman, D. et al. Optical Pooled Screens in Human Cells. Cell 179, 787–799 e717, doi:10.1016/j.cell.2019.09.016 (2019).

36 Sanchez, H. M., Lapidot, T. & Shalem, O. High-throughput optimized prime editing mediated endogenous protein tagging for pooled imaging of protein localization. bioRxiv, doi:10.1101/2024.09.16.613361 (2024).

37 Tan, M. & Le, Q. in *International conference on machine learning*. 6105-6114 (PMLR).

38 Ronneberger, O., Fischer, P. & Brox, T. in Medical image computing and computer-assisted intervention–MICCAI 2015: 18th international conference, Munich, Germany, October 5-9, 2015, proceedings, part III 18. 234–241 (Springer).

39. Courty, N., Flamary, R. & Ducoffe, M. Learning wasserstein embeddings. arXiv preprint arXiv:1710.07457 (2017).

40 Haviv, D., et al. Wasserstein wormhole: Scalable optimal transport distance with transformers. *ArXiv*, arXiv: 2404.09411 v09414 (2024).

41 Flamary, R. et al. Pot: Python optimal transport. Journal of Machine Learning Research 22, 1–8 (2021).

42 Farahmand, A. m., Szepesvári, C. & Audibert, J.-Y. in *Proceedings of the 24th international conference on Machine learning* 265–272 (Association for Computing Machinery, Corvalis, Oregon, USA, 2007).

43 Tenenbaum, J. B., de Silva, V. & Langford, J. C. A global geometric framework for nonlinear dimensionality reduction. Science 290, 2319–2323, doi:10.1126/science.290.5500.2319 (2000).

44 Santhosh Kumar, S., et al. Sequential CRISPR screening reveals partial NatB inhibition as a strategy to mitigate alpha-synuclein levels in human neurons. Sci Adv 10, eadj4767, doi:10.1126/sciadv.adj4767 (2024).

45 Santhosh Kumar, S., et al. Sequential CRISPR screening reveals partial NatB inhibition as a strategy to mitigate alpha-synuclein levels in human neurons. Science Advances 10, eadj4767 (2024).

