## Supplementary Figures for "Morphology-robust quantification of subcellular organization in complex cells"

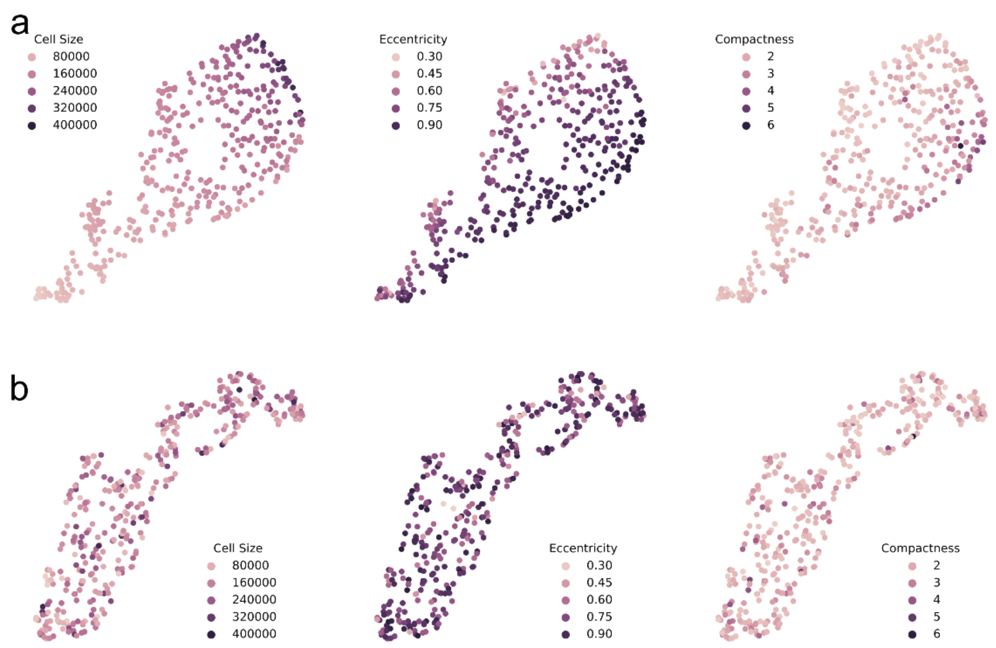


**Supplementary Figure 1. Distribution of cell morphological features in the CellAligner cell morphology and subcellular localization spaces for 387 cells from the Human Protein Atlas. (a)** UMAP of the CellAligner cell morphology space, colored by cell size, eccentricity, and compactness. **(b)** UMAP of the CellAligner-OT subcellular localization space, colored by cell size, eccentricity, and compactness.

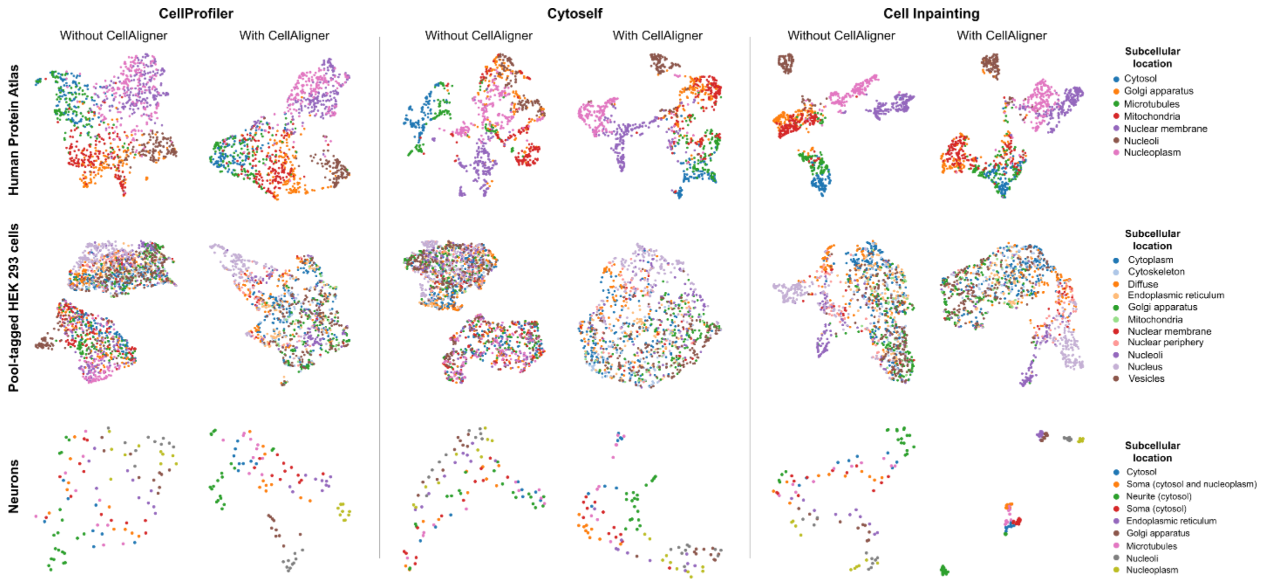


**Supplementary Figure 2. UMAP embeddings of subcellular localization spaces constructed from the outputs of various current methods.** Embeddings are shown for the three evaluation datasets (Human Protein Atlas images, pool-tagged HEK293 cells, and neurons with simulated subcellular localization patterns) and methods (CellProfiler, Cytoself, and Cell Inpainting) considered in Fig. 2 applied either to the original images or to the anchor-cell representations produced by CellAligner.

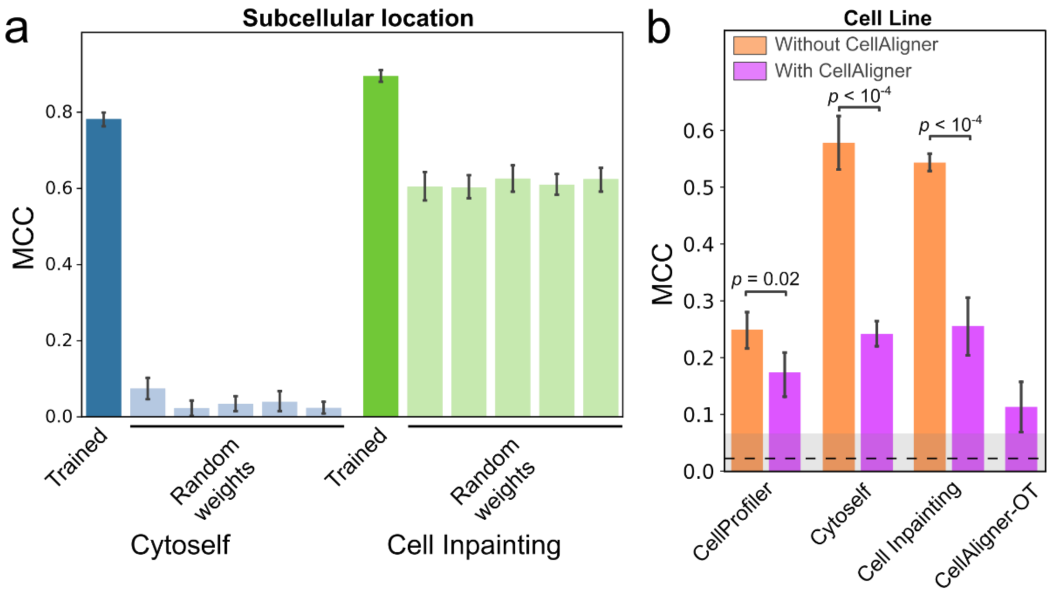


**Supplementary Figure 3. Quantitative evaluation of CellAligner performance. (a)** MCC of k-nearest neighbor classifiers trained to predict subcellular localization labels from the outputs of Cytoself and Paired Cell Inpainting, evaluated for the full pre-trained model and for randomly initialized weights without training, across five different random initializations. **(b)** MCC of k-nearest neighbor classifiers trained to predict cell-line identity from the outputs of CellAligner-OT and three subcellular localization analysis methods applied either to the original images or to the anchor-space representations generated by CellAligner. The dashed line and grey band indicate the baseline performance, defined by the average MCC ± standard deviation of a random classifier. Two-sided Wilcoxon rank-sum test p-values are shown when statistically significant ($p \leq 0.05$).

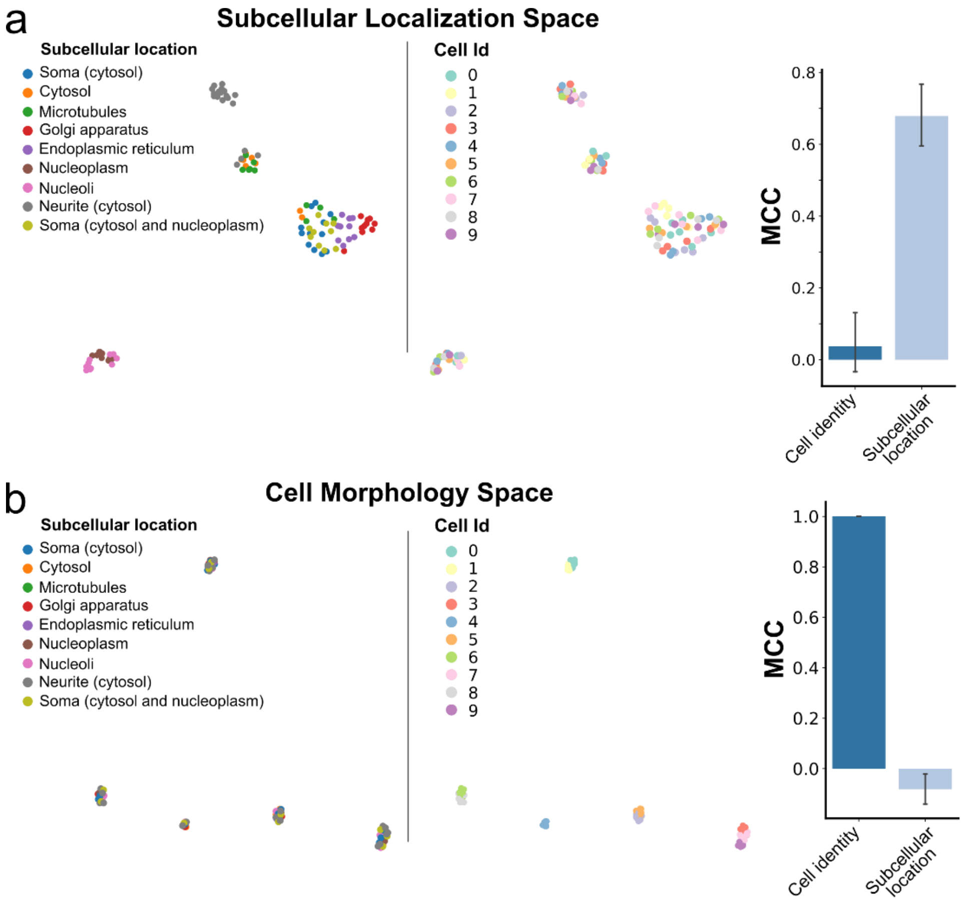


**Supplementary Figure 4. Quantitative evaluation of CellAligner on images of neurons with simulated subcellular localization patterns. (a)** UMAP embedding of the subcellular localization space generated by CellAligner-OT, colored by subcellular localization (left) and cell identity (center) labels, and MCC of a k-nearest neighbor classifier trained to predict cell identity and subcellular localization labels from this space (right). **(b)** UMAP embedding of the cell morphology space generated by CellAligner, colored by subcellular localization (left) and cell identity (center) labels, and MCC of a k-nearest neighbor classifier trained to predict cell identity and subcellular localization labels from this space (right).

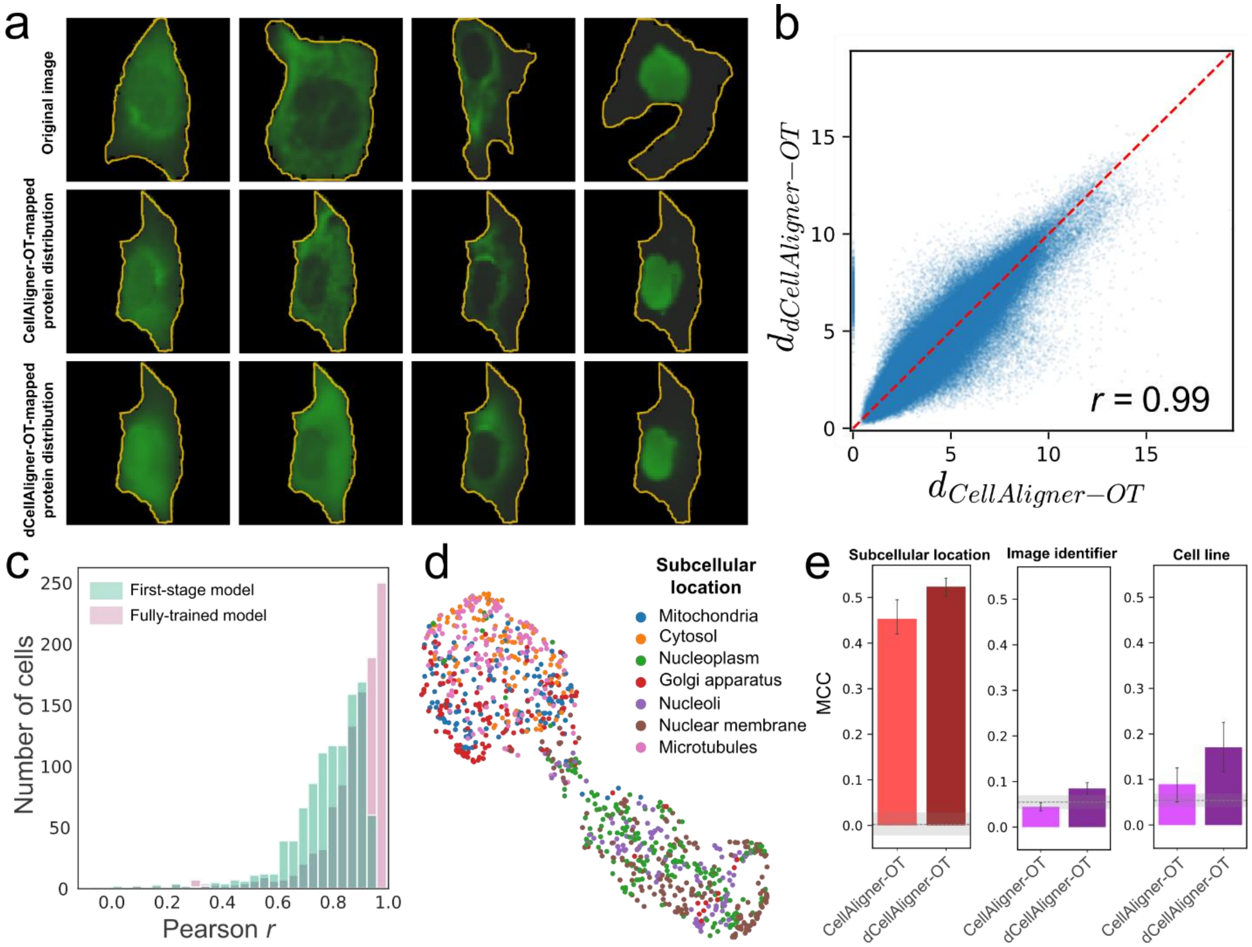


**Supplementary Figure 5. Evaluation of dCellAligner-OT on Human Protein Atlas Data.** (**a**) Examples of Human Protein Atlas cell images mapped onto a reference anchor cell using CellAligner-OT or the trained dCellAligner-OT model. (**b**) Scatter plot comparing pairwise CellAligner-OT distances with pairwise dCellAligner-OT distances in the Human Protein Atlas test dataset, for both first-stage and fully-trained dCellAligner-OT models. (**c**) Distribution of pixel-wise Pearson correlation coefficients between CellAligner-OT and dCellAligner-OT mapped protein distributions. (**d**) UMAP representation of the subcellular localization space produced by CellAligner-OT for the Human Protein Atlas test dataset, colored by ground-truth subcellular localization labels. (**e**) MCC of k-nearest neighbor classifiers trained to predict subcellular localization, image identifiers, and cel lines based on the subcellular localization space produced by dCellAligner-OT for the Human Protein Atlas test dataset. The dashed line and grey band indicate the baseline performance, defined by the average MCC ± standard deviation of a random classifier. For image-identity and cell-line-identity prediction, baseline performance was computed after conditioning on the localization label.
